# Assessing the impact of character evolution models on phylogenetic and macroevolutionary inferences from fossil data

**DOI:** 10.1101/2024.12.23.630137

**Authors:** David F. Wright, Melanie J. Hopkins

**Affiliations:** Sam Noble Oklahoma Museum of Natural History, 2401 Chautauqua Avenue, Norman, OK, 73072, USA; School of Geosciences, University of Oklahoma, 100 E Boyd Street, Norman, OK, 73019, USA; Department of Paleobiology, National Museum of Natural History, Smithsonian Institution, Washington, DC, USA; American Museum of Natural History, Division of Paleontology (Invertebrates), Central Park West at 79th St., New York, NY 10024, USA

**Author notes:** Corresponding author: David F. Wright.

**Keywords:** palaeobiology, morphology, phylogenetics, macroevolution, fossil data

## Abstract

Understanding the evolution and phylogenetic distribution of morphologic traits is fundamental to macroevolutionary research. Despite decades of major advances and key insights from molecular systematics, organismal anatomical features remain a key source of biological data for both inferring phylogenies and investigating patterns of trait evolution among fossil and extant species. In palaeobiology, morphologic characters are typically the only source of information available for reconstructing evolutionary trees. Systematists working with fossil data must make decisions regarding how morphological characters are modeled, whether they are coded as continuous or categorical, and how to address biological sources of rate variation. To determine the impact of how different models of morphological evolution influence phylogenetic inferences and downstream comparative analyses of fossil data, we compared a series of increasingly complex model configurations of character evolution to a dataset of Cambrian-Ordovician trilobites containing both discrete morphological characters and continuous traits. Model configurations vary in complexity, ranging from simple constant rate scenarios with only discrete categorical traits, to complex evolutionary models including both discrete and quantitative traits across multiple ecological partitions while accounting for multiple sources of rate variation. We compared topological distributions across model configurations by visualizing their distances in multidimensional treespace. Results indicate support for rate-variable models and partitioning characters. However, inclusion of continuous traits dramatically alters macroevolutionary inferences. Character model complexity also has a major impact on which regions of treespace are explored, suggesting a critical need to carefully model morphologic evolution even when systematists are only interested in patterns of phylogenetic relationships alone.

## 1. Introduction

Investigating patterns of anatomical character evolution is a major component of macroevolutionary biology (Simpson, 1944; Harmon et al., 2010). Character evolution models are fundamental to documenting evolutionary trends (Monnet et al., 2011; Hopkins and Lidgard, 2012), interpreting patterns of morphological disparity in the fossil record (Foote, 1997; Erwin, 2007), and testing alternative hypotheses of trait evolution using time-calibrated phylogenetic trees (Revell and Harmon, 2022). Moreover, despite decades of major advances and key insights from molecular systematics, organismal anatomical features remain a key source of biological data for both inferring phylogenies and studying patterns of trait evolution among fossil and extant species (Ronquist et al., 2012; Wright and Hillis, 2014; Bapst et al., 2018; A. Wright, 2019). In some disciplines, such as palaeobiology, anatomical characters are frequently the most common and perhaps only source of information available for reconstructing phylogenies (Smith, 1994; Wiley and Lieberman, 2011). Most phylogenies of fossil taxa have historically been inferred using parsimony methods (e.g., Smith 1994; Cobbett et al., 2007; O’Reilly et al., 2018; Sansom et al., 2018). However, over the last several years, systematists working with fossil data have increasingly begun using Bayesian hierarchical modeling approaches to simultaneously infer a tree topology, date divergences, and estimate macroevolutionary parameters (Wright, 2017a; Warnock and Wright, 2020; Wright et al., 2021; Mulvey et al., 2025). In particular, Bayesian “tip-dating” phylogenetic methods that implement the fossilized birth-death process (FBD) (Stadler, 2010; Heath et al., 2014; Gavryushkina et al., 2014; Stadler et al., 2018) are especially appealing to palaeobiologists because they can leverage both morphological and stratigraphic data from the fossil record to estimate a probability distribution of evolutionary histories (i.e., relationships and macroevolutionary dynamics) (Wright, 2017a; Paterson et al., 2019; King and Beck, 2020; Barido-Sottani et al., 2023).

Simulations show the inclusion of fossil age information under the FBD process can improve inferred tree topologies for tip-dated analyses relative to undated approaches (Barido-Sottani et al., 2020; Mongiardino Koch et al., 2021). However, the inclusion of fossil age data in tip-dating studies predominantly impacts the probability of tree shapes and diversification histories (May and Rothfels, 2023), whereas morphological character data remain the primary source of information regarding the probability of alternative topologies (Warnock and Wright, 2020; Wright et al., 2021). Since appropriately modeling fossil age data can improve phylogenetic inferences (Barido-Sottani et al., 2020), it seems even more prudent to determine the extent to which inferences may also be influenced by different approaches to handling assumptions about character evolution, especially in empirical studies where the tree topology is often the parameter of greatest interest. Unfortunately, when one considers the complexity of information captured by different kinds of morphological characters and the vast number of ways in which they could be modeled, determining how to appropriately model character evolution remains a challenging task.

Discrete characters comprise the majority of morphological matrices in the literature (Wright, 2019; Mulvey et al., 2025). The simplest implementation of a discrete character evolution model is the Mk model of Lewis (2001), which assumes characters are conditionally independent, evolve at uniform rates, and have symmetrical transition frequencies between character states. Although the Mk model has been shown to outperform parsimony under a variety of scenarios (Wright and Hillis, 2014), many empirical matrices exhibit dramatic heterogeneity in morphologic rates among characters, across lineages, and through time (Lloyd et al., 2012; Hopkins and Smith, 2015; Wright, 2017b). Consequently, a variety of approaches have been developed to relax assumptions about character evolution and accommodate different sources of rate variation among characters and lineages (Wagner and Marcot, 2010; Drummond and Stadler, 2016; Wright et al., 2016). For example, rate variation among characters is often modeled using a parametric distribution, such as a gamma or lognormal (Wagner, 2012; Harrison and Larsson, 2015; Wright, 2019). The variance of the rate distribution, which determines its shape, is a single parameter estimated from the data. Because per-character rates are independent draws from the same distribution, character evolution is implicitly modeled as having a common generating process but with variable rates. However, not all characters might be expected to evolve according to the same evolutionary “rules” or generating mechanisms. In such instances, characters may be partitioned into categorically different groups, each having unique parameters to describe their dynamics. For example, different anatomical regions may vary in morphologic rates or involve developmentally-linked correlated character changes (Clark and Middleton, 2008; Wagner, 2018). Alternatively, characters with ecological or functional significance may evolve at higher rates than non-ecologic traits due to elevated selective pressures that fluctuate in direction over time (Blomberg et al. 2013).

Another source of morphologic data comes from continuous characters, which are commonly modeled using diffusion processes such as Brownian motion (BM) or related models (Felsenstein, 1988). Although less frequently used to infer phylogenies, continuous characters may be used to infer both tree topology (Parins-Fukuchi, 2018a; Zhang et al., 2024) and divergence times (Álvarez-Carretero et al., 2019). When modeled directly, continuous characters have been shown to contain at least as much, if not greater, phylogenetic signal than discrete characters (Parins-Fukuchi, 2018b). However, in many empirical matrices, continuous variation is binned into categories and treated as a discrete character (e.g., Rae, 1998; Thiele, 1993; for examples see Gahn and Kammer, 2002; Wright and Stigall, 2013, 2014; Ausich et al., 2015), and the extent to which this process of discretization results in the loss of phylogenetic information is unclear (Rae, 1998).

The wide variety of ways to model morphological evolution leads to a large number of possible model configurations, and computational tractability prevents evaluation of all possible candidate models using conventional model selection techniques, such as Bayes Factors (Wright et al., 2021). Unlike substitution models for molecular sequence data, few frameworks exist for determining which character model configuration provides a best fit to morphologic data in advance (Lanfear et al., 2017). Remarkably, the practice of fitting molecular substitution models to sequence data prior to phylogeny reconstruction has previously been argued to be unnecessary if the purpose is solely to infer the tree topology or conduct ancestral state reconstruction, as both simulated and empirical results tend to be similar to the most parameter-rich substitution model regardless of the generating process or best fit to empirical data (Abadi et al., 2019). If these results extend to morphological matrices of fossil taxa, then the time-consuming steps involved in model selection may be similarly unnecessary, thereby liberating palaeobiologists from the need to carry out a series of difficult and computationally expensive analyses prior to phylogeny reconstruction. In this study, we investigate this question empirically using a morphological dataset of olenid trilobites, a major clade of extinct arthropods. Specifically, we (1.) compare a series of increasingly complex character model configurations to these data, (2.) conduct model selection to rank configurations by their relative fit, and (3.) compare the evolutionary implications associated with inferences made under each configuration. The goal of this study is to assess the degree to which different character model configurations alone may impact phylogenetic and macroevolutionary inferences in empirical analyses of fossil data.

## 2. Materials and methods

### (a) Data

We focus our analyses on trilobites, a species-rich and morphologically diverse clade of Palaeozoic arthropods, as a model system for evaluating character evolution models in phylogenetic palaeobiology. We selected a morphological character matrix from a previously published parsimony-based analysis of the Cambrian – Ordovician trilobite family Olenidae that sampled both discrete and continuous characters (Monti and Confalonieri, 2019), and subsequently updated and modified the character matrix in two ways. First, we coded two additional species from the subfamily Balnibarbiinae, which was unrepresented in the original matrix (*Balnibarbi pulvurea* and *Cloacaspis senilis*, see Fortey 1974; Hopkins, 2019). Next, we noted the original matrix contained a substantial number of cells with missing entries for continuous characters. Since we are primarily interested in how different character models may impact downstream inferences rather than reevaluating previous authors’ results, for computational tractability (Barido-Sotanni et al., 2023), we removed a combination of taxa and continuous characters in an effort to retain the maximum number of entries with no missing cells. However, all discrete morphologic characters from the original matrix were retained. The final dataset includes 38 species, 62 discrete characters, and 12 continuous traits, which is a relatively large character matrix for fossil marine invertebrates (Barido-Sottani et al., 2020), and we believe likely representative of many published morphologic datasets of similar size and taxonomic scope. Stratigraphic occurrence data for each fossil species were compiled from the literature and species were assigned ages corresponding to geologic intervals bounding their first appearance intervals using Gradstein et al. (2012). The character matrix and data are provided in the Supplemental Information.

### (b) Character evolution models

We compared five morphologic model configurations that range from simple to complex, with more parameter-rich models accounting for increasingly more biologically complex evolutionary dynamics (Figure 1). In the first three model configurations, all morphologic characters analyzed are discrete traits (e.g., “absence” or “presence” of a character), whereas the two most complex configurations also include continuous characters and implement separate character evolution models for discrete versus quantitative traits.

**Figure 1.**
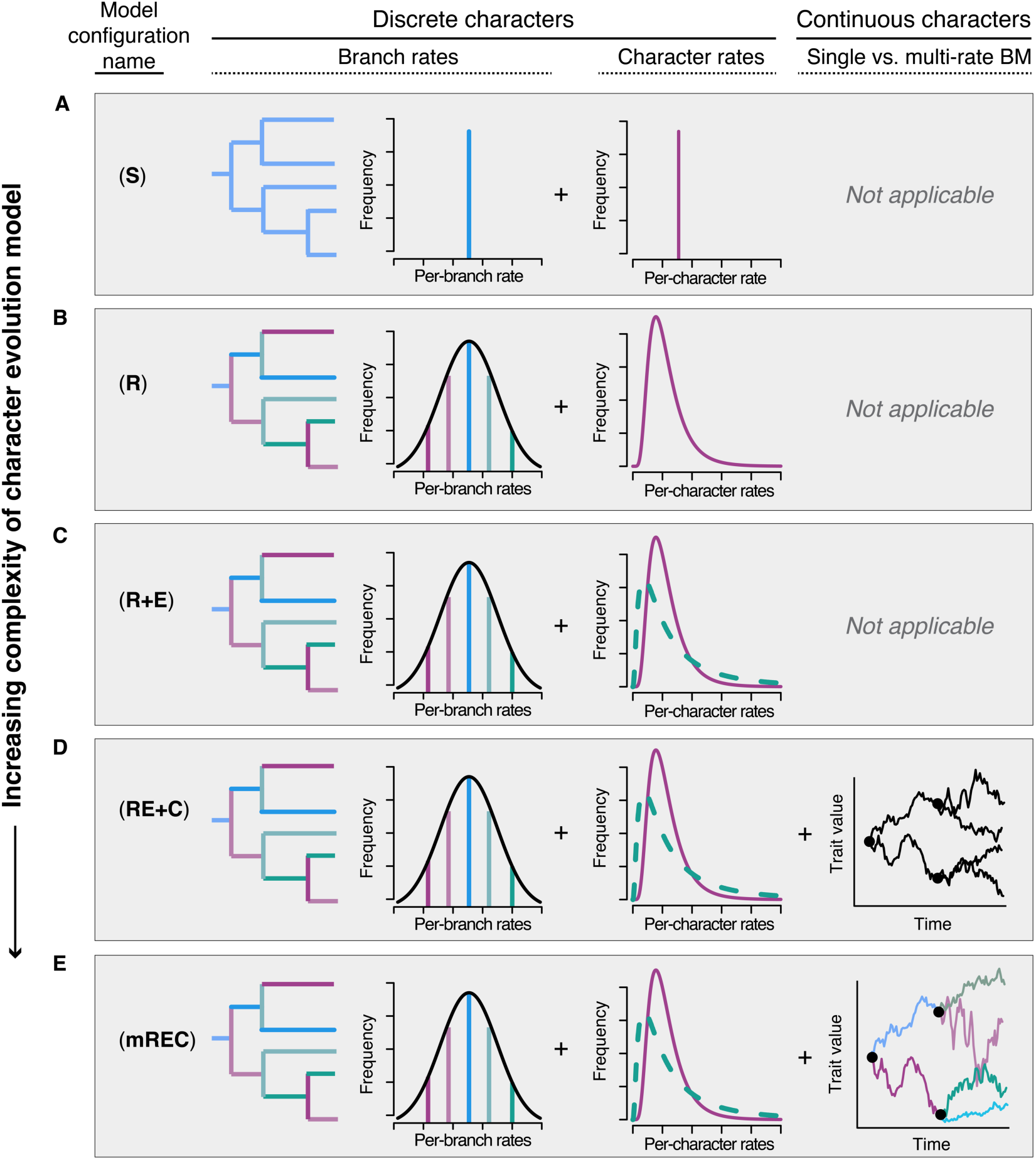
Character evolution model configurations. Model configurations correspond to a range of simple to increasingly complex and parameter-rich models of morphological evolution; (**A**) model **S**, strict morphological clock of discrete morphological data with equal rates of change among characters; (**B**) model **R**, allows for rate variation among characters and across phylogeny; (**C**) model **R+E**, builds on model **R** with traits identified as ecologic vs. non-ecologic characters assigned to different character partitions; (**D**) model **RE+C**, builds on **R+E** but with continuous characters evolving according to a single-rate Brownian motion process; (**E**) model **mREC**, modifies model **RE+C** such that continuous characters evolve according to a multi-rate Brownian motion process.

The simplest model configuration (**S**) corresponds to a constant rates model assuming a “strict” morphological clock and an equal rates implementation of the Mk model of discrete character evolution (Lewis, 2001). In contrast, the “relaxed” rates model (**R**) allows for discrete character rates to vary among characters and through time. Specifically, rates vary among characters following a lognormal distribution (Wagner, 2012), and across the phylogeny according to an uncorrelated lognormal clock model (Heled and Drummond, 2012). Next, the configuration of the **R+E** model modifies the **R** model configuration by partitioning “ecologic” characters and estimating separate lognormal rate distributions for traits with presumed ecological significance vs. non-ecological traits (**R+E**). Ecological significance was defined as any character relevant to sensory systems (e.g. size and placement of eyes) or behavior (e.g. molting or defense). A total of 13 characters (20%) were assigned to the “ecological” partition. Next, the **RE+C** model configuration uses the **R+E** configuration for discrete traits but includes continuous characters and models them as quantitative traits evolving according to a single rate Brownian motion process (Felsenstein, 1985). Finally, the most complex model implements a multivariate BM model for continuous traits (**mREC**), which accommodates rate variation among quantitative characters and accounts for evolutionary integration among correlated traits (Caetano and Harmon, 2019).

### (c) Phylogenetic and macroevolutionary methods

All analyses were conducted using “tip-dating” Bayesian phylogenetic methods incorporating the fossilized birth-death (FBD) process (Stadler, 2010; Heath et al., 2014; Wright, 2017a). The tree prior is an important component of Bayesian phylogenetic inference, especially for datasets including many (or only) fossil taxa (Barido-Sottani et al., 2020; Warnock and Wright, 2020).

However, fossil occurrence times are not just priors for node ages and/or tip occurrences, they are also data, and changing our assumptions about how the tree prior is handled may inadvertently prevent us from discerning which differences are due to treatment of character data vs. tree priors (May and Rothfels, 2023). Thus, to focus solely on the impact of alternative character evolution models, we applied a simple time-homogeneous, sampled ancestor implementation of the FBD process across all model configurations.

The relative fit of alternative model configurations were assessed using Bayes factors (BF) following the stepwise model selection process outlined by Wright et al. (2021) for palaeontological datasets, a common practice in phylogenetic palaeobiology. Calculating Bayes factors involves estimating and comparing marginal likelihoods for the set of competed models, where the marginal likelihood is the probability of the data integrated over a model’s parameter space. Marginal likelihood estimation was conducted using Stepping-Stone (SS) sampling in RevBayes (Xie et al., 2011; Höhna et al., 2016) following the algorithm and procedures for generating power posteriors using SS sampling described in Höhna et al. (2024). Model-fitting using SS sampling estimates marginal likelihoods by iteratively sampling parameter space from the prior to the posterior, which approximates the sum of the probability of the data across all possible values of parameter space. Once marginal likelihoods were obtained for each model configuration, Bayes factors were calculated and sequentially compared among each successively more complex pair of models. To facilitate comparison of fit across model configurations with different data types (i.e., discrete-only vs. discrete and continuous traits), BFs for models with only discrete data were compared and evaluated separately from models including discrete and continuous traits. Interpretation of BFs follow guidelines of the natural log posterior probability scale in Kass and Raftery (1995).

In addition to comparing the relative fit among models, we also wanted to determine the extent to which different model configurations might lead to different phylogenetic hypotheses and/or macroevolutionary inferences. To compare the posterior distributions for tree topologies and macroevolutionary parameters inferred under different models, we conducted Markov-chain Monte Carlo (MCMC) simulation in RevBayes (Höhna et al., 2016) for each of the character model configurations (Figure 1). Macroevolutionary parameters of interest include FBD parameters for speciation rate, extinction rate, and sampling rate, as well as the number of sampled ancestors in each tree of the posterior distribution. To evaluate whether character models impact node ages and/or branch durations, we also calculated the minimum implied gap (MIG) (Wills, 1999; O’Connor and Willis, 2007), which is equal to the sum of the ghost ranges implied by a timescaled phylogeny when terminal stratigraphic ranges are excluded. To determine whether clade support values were influenced by character model configurations, we inferred a Maximum Clade Credibility (MCC) tree for each character model configuration and compared their posterior probabilities of clades. Finally, to compare the samples of trees inferred by different character model configurations, we mapped the posterior distribution of phylogenies inferred by each model configuration into a “treespace” (Hillis et al., 2005; Wright and Lloyd, 2020; Pates et al., 2022). Treespaces were constructed via multivariate ordination of pairwise dissimilarity matrices using Principal Coordinate Analysis (PCoA). Pairwise dissimilarity among trees was calculated using a variety of tree distance metrics, including the simple Robinson-Foulds (RF) distance, which measures the number of splits that differ between trees (Robinson and Foulds, 1981), as well as information theoretic methods (Smith, 2020) and quartet distances (Estabrook et al., 1985).

Convergence of SS sampling and MCMC results were evaluated using standard approaches to MCMC diagnostics, including visual inspection of trace plots, comparing independent runs, and checking estimated sample sizes. All treespace analyses were performed in R using original scripts and functions from the ape package (v. 5.7; Paradis and Schliep, 2019), the RF.dist function from the *phangorn* package (v. 2.10.0; Schliep, 2011), the ClusteringInfoDist function from the *TreeDist* package (v. 2.9.2; Smith, 2022), the TQDist function from the *Quartet* package (v. 1.2.5; Sand et al., 2014; Smith 2019), and the cmdscale function from the *stats* (v. 4.2.1) base R package. Additional information about treespace construction, pairwise dissimilarity metrics, and custom R scripts to carry out treespace analyses are provided in the Dryad Digital Repository [link].

## 3. Results

The log marginal likelihood values for character model configurations and BFs for each sequential model comparison and data types are shown in Table 1. Bayes Factors among competed models generally indicate increased support for increasingly complex character models with additional layers of biological complexity. However, the explanatory data available limits this trend, as BFs naturally penalize more parameter-rich models (Kass and Raftery, 1995), and our results decisively indicate the most parameter-rich model is not necessarily supported compared to a simpler model configuration.

**Table 1.**
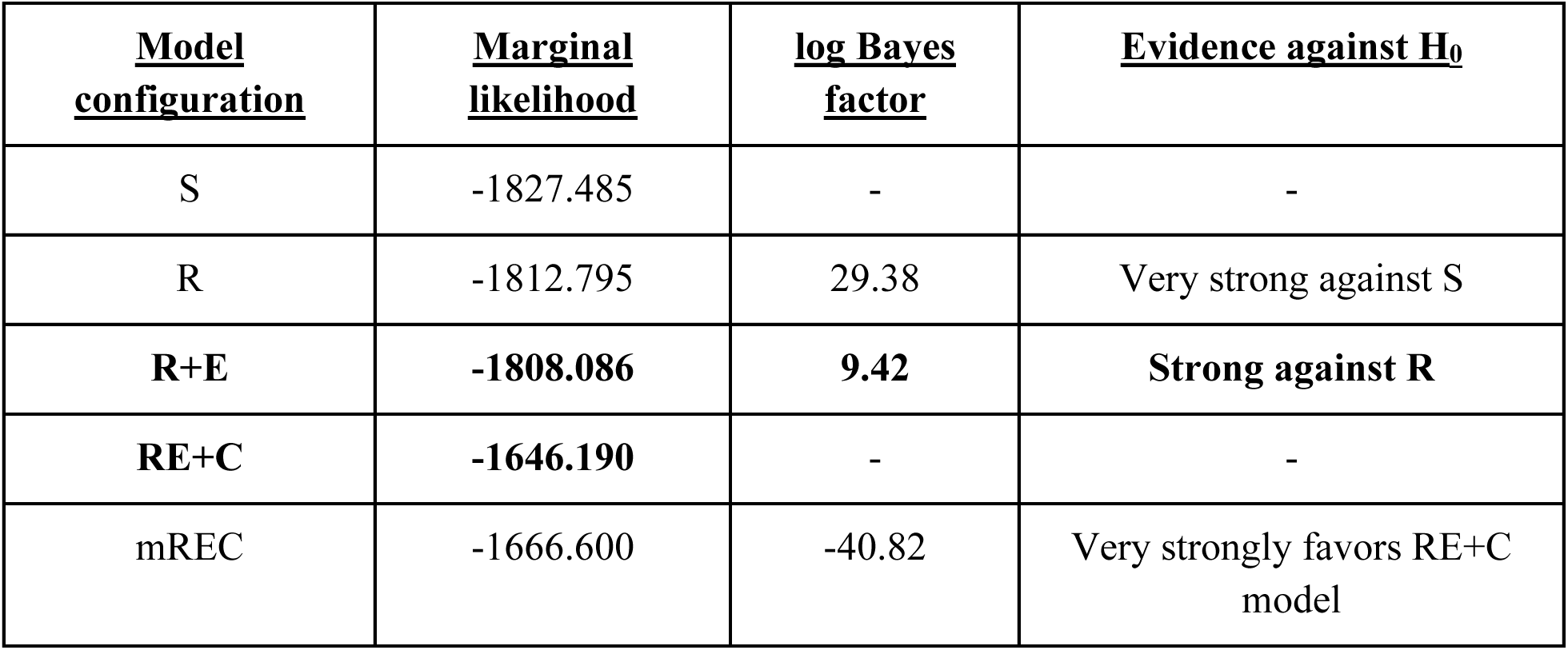
Results of model selection comparison for the two types of character sets. The log Bayes factor scale and interpretation follows Kass and Raftery (1995). R+E is favored for the discrete character data models, whereas RE+C is favored for the discrete + continuous data models.

Among models analyzing only discrete traits, BFs indicate the **R+E** configuration was the best-fitting model, which accounted for multiple patterns of rate variation and partitioned characters into categories comprising “ecologic” vs. non-ecologic features of the trilobite body plan. The inclusion of continuous traits dramatically increased log marginal likelihood values compared to model configurations with only discrete traits. Among the two model configurations analyzing both discrete and continuous traits, BFs indicate the **RE+C** model was favored over **mREC**, and had the larger log marginal likelihood value and decisive BF support.

The posterior distributions of macroevolutionary and sampling parameters estimated under different character evolution models show considerable variation (Figure 2). For example, speciation and extinction rates were inferred to be considerably higher for rate-variable models, whereas fossil sampling rates are estimated to be lower on average compared to a constant rate model. The best-fitting character model, **RE+C**, has the lowest mean fossil sampling rate among models. According to MIG values, models incorporating continuous trait evolution imply longer durations of “ghost” lineages than other rate-variable models that sample only discrete characters, and on average contain fewer sampled ancestors per tree than models without continuous trait data (Figure 2). Maximum clade credibility (MCC) trees inferred across model configurations show substantial variation in posterior probability values (Figure 2). Since MCC trees are fully bifurcating, support values for clades corresponding to their *N*-1 internal nodes can be compared. Interestingly, the **S** model, which has the lowest marginal likelihood and BF support among models with only discrete characters (Table 1), has the highest mean posterior probability per clade across the 37 internal nodes. However, we find no statistical differences in clade posterior probabilities recovered in MCC trees across model configurations (one-way anova: *F* = 1.17, *df* = 4, *p* = 0.326). The MCC trees inferred under each character model are presented in Appendix S1, S19-23.

**Figure 2.**
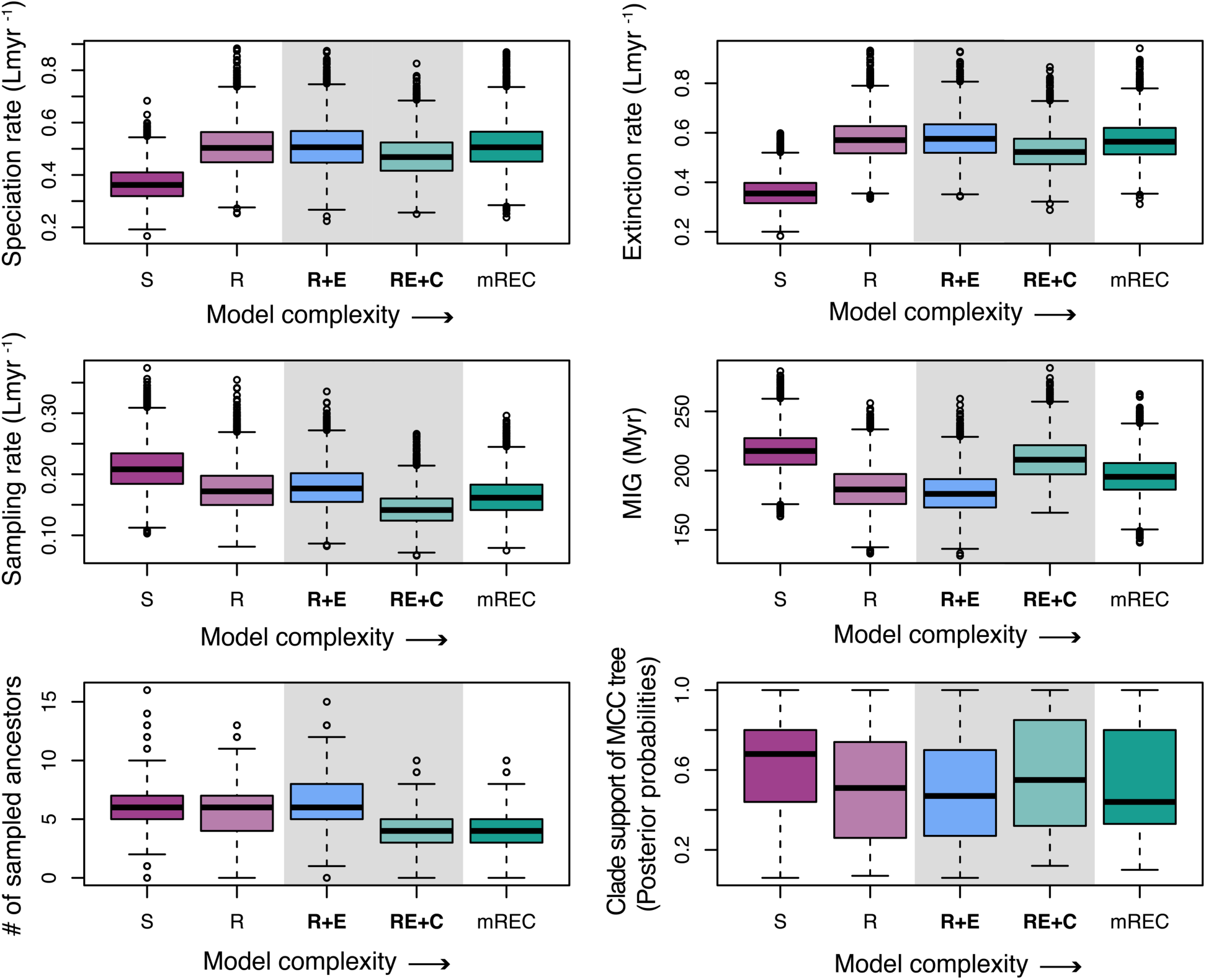
Impact of character models on macroevolutionary and phylogenetic inferences. Except for clade support values (lower right), all boxplots show variation in parameter estimates across the posterior distribution resulting from MCMC analysis of each model configuration. Speciation, extinction, and sampling are in units of events per-lineage-million-years (Lmyr ^-1^). Clade support indicates variation in tree-wide posterior probability of clades in the MCC tree. Abbreviations for model configurations as for Figure 1. MIG indicates the minimum implied gaps. Model configurations with the highest support for the discrete and discrete + continuous character sets are indicated in **bold** with a gray background.

Results of treespace visualization and analysis of tree-to-tree properties (e.g., tree balance) were strongly concordant with one another across a variety of alternative distance metrics (see Appendix S1). For simplicity, we focus our main results on pairwise distances calculated using the simple RF metric. Scree plots, projections of tree space onto other principal coordinates, and comparisons with tree spaces based on other tree samples and other distance measures are reported in the Appendix S1.

A visualization of treespace projected along the first two PCoA eigenvectors is shown in Figure 3. Although trees inferred under different character model configurations show some overlap in the two-dimensional space of PCoA axes 1 and 2, the treespace is highly structured with respect to the density of points and their locations along PCO1, which shows a striking correspondence with character model complexity (Figures 1 and 3). Moreover, the MCC trees inferred under different models are highly dispersed throughout the treespace, and clusters of trees form “islands” within treespace such that more complex model configurations are almost non-overlapping with the simplest character evolution model. On average, the posterior distributions of trees for model configurations including continuous characters contain trees that are less balanced, and the least complex model returned a larger proportion of balanced trees compared to other model configurations (Appendix S1).

**Figure 3.**
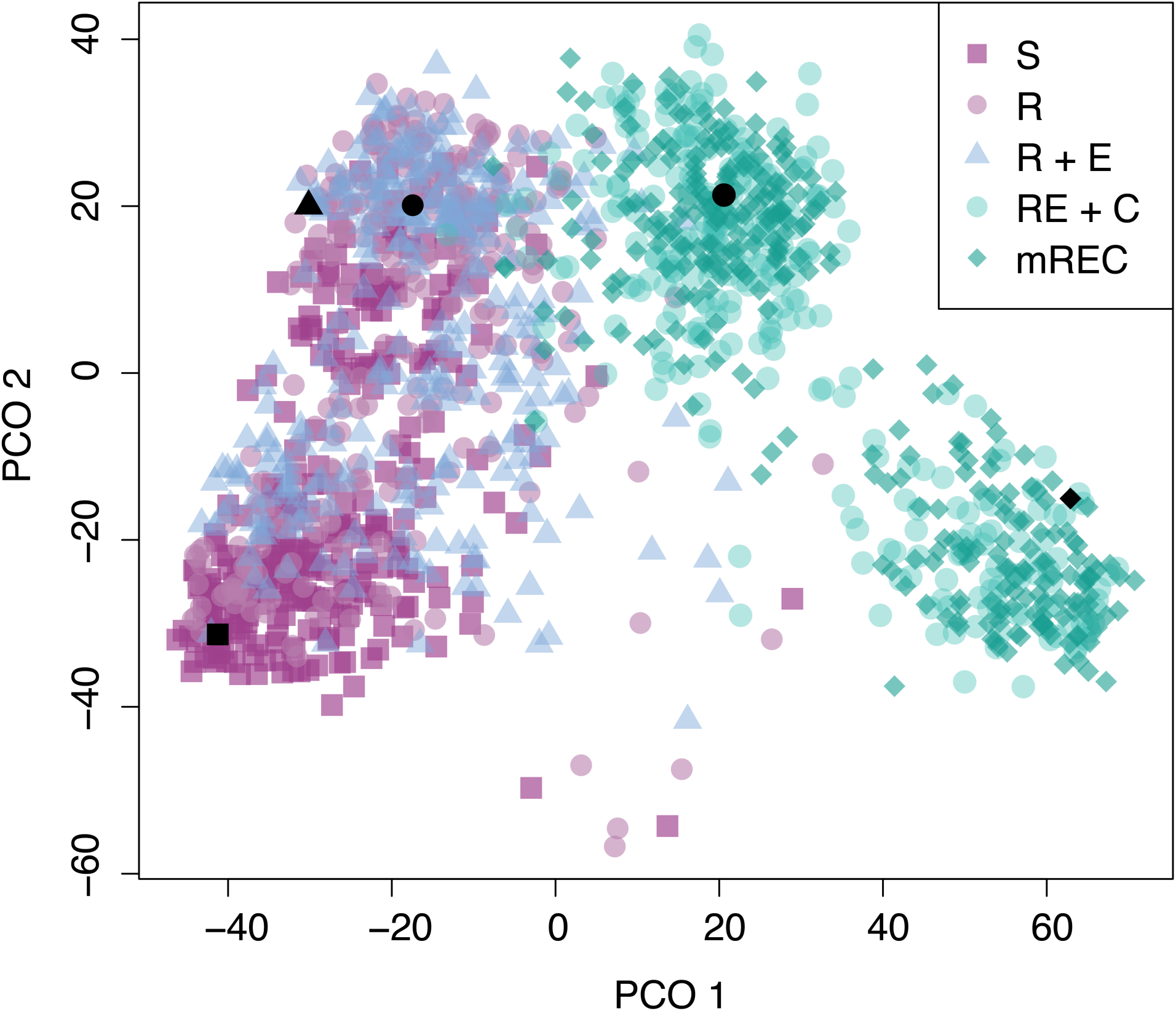
Tree space of 300 randomly selected trees from each posterior distribution. The tree space is a PCoA of a distance matrix based on RF distances between all pairs of trees; shown here is a projection of tree space onto the first two principal coordinates (PCO 1 and PCO 2). Black shapes correspond to the MCC trees (Appendix S1, S19-23) for each model configuration. See Figure 1 caption for model configuration abbreviations.

## 4. Discussion

### (a) Reconstructing evolutionary history: a place where all the different kinds of truths fit together

In comparative biology, knowledge of evolutionary relationships among species makes predictions regarding patterns of character change (Felsenstein, 1985; Harmon, 2019; Soul and Wright, 2021). But in morphological systematics, especially phylogenetic palaeobiology, character evolution models are themselves used to infer phylogenetic hypotheses—a seemingly catch-22 situation. As a further complication, we note even simple models of character evolution across a phylogeny, such as constant rate Brownian motion, can generate highly complex patterns of trait evolution when speciation and extinction rates vary over a clade’s lifetime (Foote, 1996; O’Meara et al., 2006). However, Bayesian phylogenetic methods naturally present a solution to these apparent dilemmas by providing a framework “where all the different kinds of truths fit together” (see Vonnegut, 1959 for a literary use of this concept). In other words, phylogenetic inferences cannot be made in a vacuum. Indeed, reconstructing evolutionary relationships is just one of the necessary ingredients for reconstructing a clade’s *evolutionary history* (i.e., tree topology, divergences, speciation and extinction rates, etc.). Using this framework, all data concerning phenotypes, fossil occurrences, and for clades with extant taxa, molecular sequences, jointly make contributions to estimating relevant evolutionary parameters and determining the probability distribution of a clade’s evolutionary history (Wagner and Marcot, 2010).

In this study, we focused our attention on how our assumptions about morphologic evolution, reflected in character models and interpretation of their parameters, may impact our understanding of phylogenetic relationships and macroevolutionary inferences. All character evolution models are wrong. Some are useful. And some are more wrong (or more useful) than others. To determine a best-fitting model, we used a stepwise approach to model selection commonly used in phylogenetic palaeobiology to statistically compare an increasingly complex series of character models (Wright et al., 2021). In our empirical example of a family of Cambrian–Ordovician trilobites, the best-fitting character evolution model is consistent with highly heterogeneous rates of morphologic evolution among characters, across the phylogeny, and between fundamentally different types of morphologic characters.

### (b) Impacts of character partitions and inclusion of continuous trait data

There has been conflicting evidence in the literature concerning whether biologically-based character partitions impact phylogenetic results (Clarke and Middleton, 2008; Porto et al., 2021; Wright et al., 2021; Casali et al., 2022). Discussion of biologically-based partitions (e.g., based on anatomical region, ecology, developmental features, etc.) is distinct from character partitions based on the number of states (Mulvey et al., 2024), which is an important but separate issue concerning how morphologic information is coded into characters and character states (Khakurel et al., 2024), and may (or may not) involve unobserved states (Tarasov, 2019).

The majority of models we analyzed partitioned the discrete morphologic characters into ecological vs. non-ecological categories and assigned them independent lognormal distributions to account for rate variation among characters. Interestingly, trees inferred under the simplest model including character partitions (**R+E**) show substantial overlap in treespace with simpler models (**S**, **R**) (Figure 3). However, Bayes factors strongly and consistently supported models including ecology-based character partitions. Aside from how partitions may impact tree topology, we wondered if the data may be pointing to an evolutionary explanation for these results. After all, the characters in our partitions differ with respect to whether or not they have a presumed functional or ecologic significance, and basic comparative biology predicts such traits may be more labile than non-ecologic traits (Blomberg et al., 2003). Recall the mean of a rate distribution is set equal to one in order for relative rates to be interpretable (e.g., Yang, 1994; Revell and Harmon, 2024), so the shape of the distribution describing rate variation among characters is often determined by a single parameter describing its variance, which is estimated from data. In Figure 4, we show the marginal distributions estimated for the rate variance parameters estimated for ecologic vs. non-ecologic character partitions under the best-fitting model. Remarkably, the distribution of rate variance parameters differs significantly between partitions (Kolmogorov-Smirnov test, *D* = 0.467, *p* < 0.001), with values estimated for characters in the ecologic partition to be larger than and outside the range of estimated values for non-ecologic characters. This suggests ecologic traits in olenid trilobites may have evolved more rapidly than their general body plan characters, which confirms biological intuition and perhaps explains why character partitions were supported by Bayes Factors.

**Figure 4.**
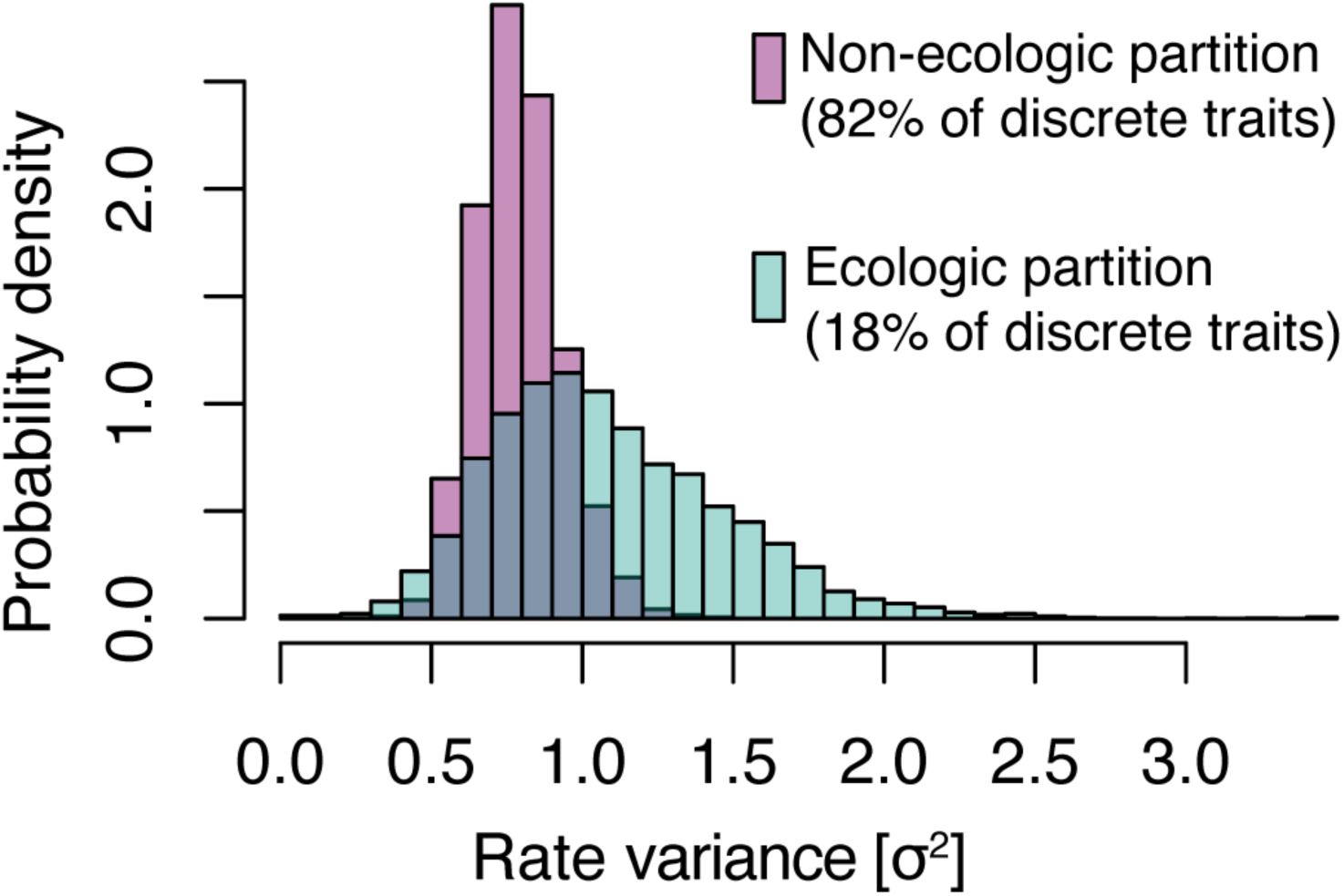
Marginal distributions of the rate variance parameters estimated for ecologic vs. non-ecologic discrete character partitions under the best-fitting character model configuration.

Although results spanning different data types cannot be directly compared via Bayes Factors, it is interesting to note that modeling continuous characters as quantitative traits dramatically increased the log marginal likelihood values (Table 1). However, the inclusion of continuous traits impacted both the macroevolutionary parameters estimated and tree topologies inferred (Figures 2-3). On average, phylogenies estimated under models including continuous traits inferred fewer sampled ancestors per tree. And perhaps for related reasons, they also have larger MIG values, indicating longer, unsampled “ghost lineage” durations across trees. We note that similar differences in node age distributions and/or branch durations have been shown to have a significant impact on downstream comparative analyses of fossil data (Bapst, 2014; Bapst et al., 2016; Bapst and Hopkins, 2017; Soul and Wright, 2019), and may be at least as influential as the overall tree topology (Soul and Friedman, 2015). Because we are dealing with empirical data, we cannot say with certainty which model (or inference) reflects the “truth”. With respect to modeling continuous traits directly (Parins-Fukuchi, 2018a), discretizing continuous variation into discrete traits (Rae, 1988; Thiele, 1993), or excluding them entirely (Pimentel and Riggins, 1987), our intuition that it is best to include as much character data and information as possible in phylogenetic analyses, provided that data reliably reflect the researcher’s understanding of their study system. Our results may be interpreted as potentially corroborating other studies highlighting use of continuous traits to estimate phylogenies and/or divergence times (Parins-Fukuchi, 2018a; Álvarez-Carretero et al., 2019). However, we caution researchers that decisions made about characters when constructing a phylogenetic character matrix may have far-reaching consequences beyond inferring relationships alone (Khakurel et al., 2024).

### (c) Character models influence the probability of tree topologies and the exploration of treespace

In addition to their impacts on macroevolutionary inferences, the character models we analyzed had a striking influence on which regions of treespace were explored during MCMC analysis (Figure 3). Regardless of whether we randomly sample 300 trees or select the 300 trees with the highest probabilities from each model’s posterior, the treespaces show similar visualizations and properties (Appendix S1). These patterns are persistent across treespaces constructed using a variety of alternative metrics to calculate pairwise distances between trees (Appendix S1), and we are unaware of any bias that could lead to our treespace ordination to show a correspondence between PCoA axes and character model complexity. Thus, a more complex character model configuration may not solely estimate more evolutionary parameters than a simpler model, it may also influence which tree topologies are considered more probable than others. As these are empirical data, we cannot know the “true” tree, but we emphasize our results indicate that our assumptions about character evolution make a difference when it comes to estimating tree topologies. Again, these results highlight how phylogenetic inference cannot be made in a vacuum (Wagner and Marcot, 2010; Wright et al., 2016).

Although the patterns in our treespace visualizations were robust to choice of tree distance metrics (Appendix S1), treespaces are complex and multidimensional. Ordination methods can cause distortion in the observed distances when projected in lower dimensional space, and it is possible that variation is spread across many axes (Hillis et al., 2005; Wright and Lloyd, 2020). To determine the fidelity of distances represented in our treespaces and assess the impact of dimension reduction, we selected two reference trees from the **S** and **mREC** models (i.e., simplest to most complex models) and compared the relationship between RF distances with Euclidean distance in the ordination using increasingly reduced sets of axes (Figure 5). In general, trees that are closer to the reference trees in Euclidean space have low RF distances, whereas trees that are further away from the reference trees have higher RF distances. We find a positive correlation between RF distances between trees and their Euclidean distances, even when only a few axes are evaluated (Figure 5). Further, similar positive correlations were found using other tree distance metrics and were not sensitive to whether treespaces were constructed using randomly selected trees or trees with the highest posterior probabilities (Appendix S1).

**Figure 5.**
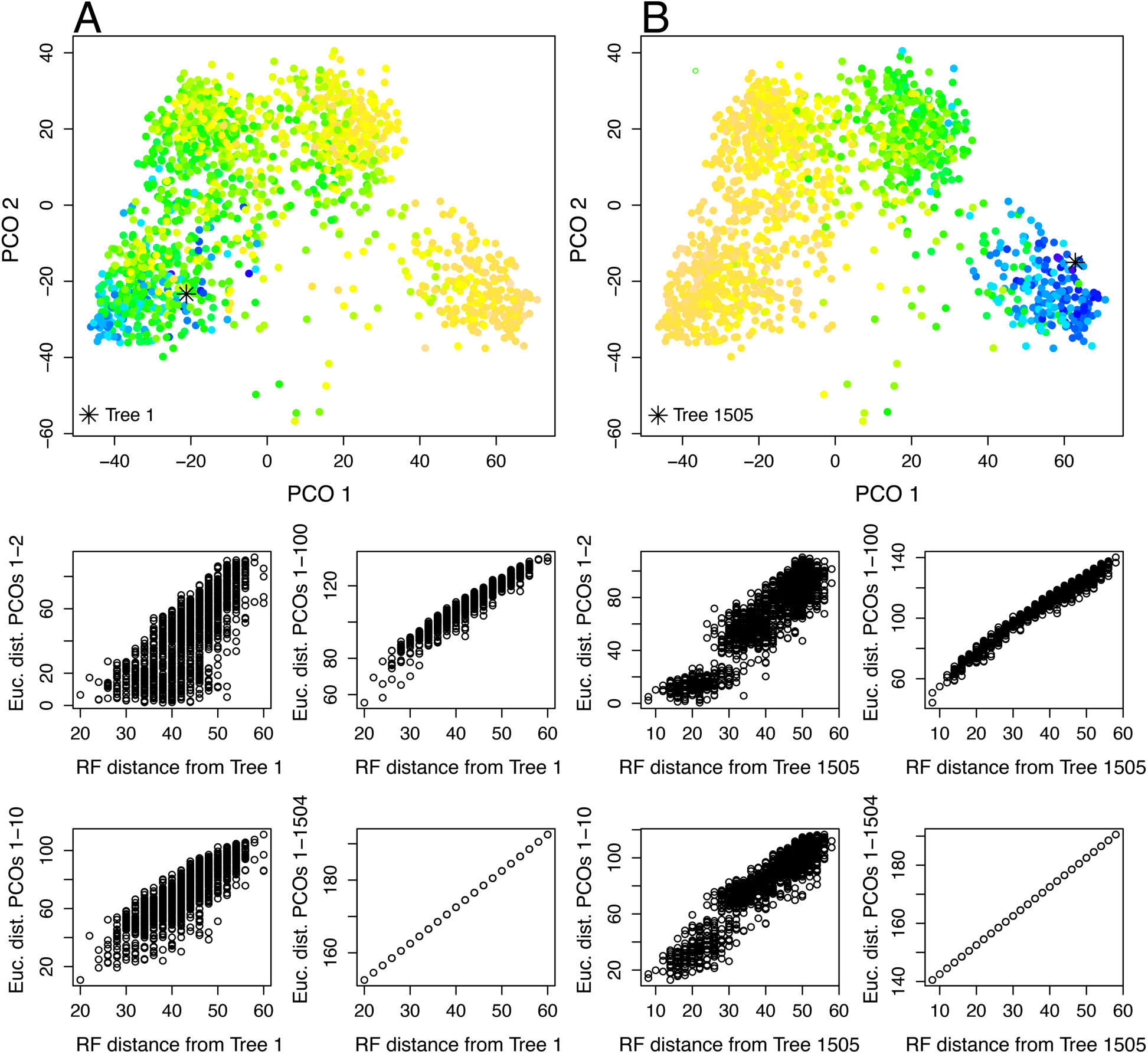
Assessment of the fidelity of projected pairwise distances and calculated pairwise distances based on the RF metric. (A) Tree space color coded by pairwise RF distance from a single randomly selected tree from the **S** model configuration’s posterior distribution (Tree 1). (B) Tree space color coded by pairwise RF distance from the MCC tree from the **mREC** model configuration’s posterior distribution (Tree 1505). Cooler colors indicate smaller RF distance; warmer colors indicate larger RF distances; lower panels show Euclidean distances calculated from subsets of PCoA scores plotted against original RF distance.

To the extent that different character models influence tree topology probabilities such that they preferentially explore different regions of treespace, we strongly recommend researchers carefully consider how they model morphological evolution and engage with model-fitting procedures even if their primary interest is solely to infer the phylogenetic relationships of their favorite clade.

### (d) Limitations, recommendations, and future directions

We evaluated only a subset of possible model configurations, and it is interesting to speculate how outcomes may vary if different decisions were made regarding the initial model choices. For example, one could have compared a model equivalent to the simplistic “strict” rates configuration (model **S**) for discrete characters, but also include the quantitative trait data and model them according to a Brownian motion process. Of course, it is neither computationally feasible nor desirable to estimate marginal likelihoods and/or compare BFs across all possible model configurations, so we chose a subset of five character evolution models we believed may represent an alternative set of plausible evolutionary dynamics for our study system. Since it is impossible to evaluate all possible models, we recommend researchers choose a subset of models that enable them to test specific evolutionary hypotheses about their clade. For example, previous studies found evidence for heterogenous per-branch rates amongst character types in trilobites (Paterson et al., 2019; Hopkins, 2017), which may be hypothesized to require a more parameter-rich model to adequately capture their evolutionary dynamics. Similarly, trilobites show considerable anatomical variation with respect to continuous traits among taxa and through time (Hopkins, 2013; Martin et al., 2023), and numerous hypotheses can be found in the literature linking aspects of trilobite morphology with species ecology (Fortey 1990; Fortey and Owens 1999; Fortey, 2014; Hopkins, 2014). Thus, the sample of model configurations we evaluated in this study are not only commonly applied to other palaeontological datasets in the literature (Wright et al., 2021; Mulvey et al., 2025), they are also sensible models to compare for olenid trilobites.

Although Bayesian model selection using SS sampling is a popular approach, it is not without several shortcomings. One possible concern of the stepwise model selection approach surrounds the order in which more complex models are evaluated, which may inadvertently bias which model is ultimately favored (Wright et al., 2021). A promising alternative to the stepwise approach to Bayesian model selection is reversible-jump MCMC (rjMCMC) (Green, 1995). In a Bayesian context, rjMCMC places priors on competing models and treats them as random variables, where models are sampled by MCMC according to their posterior probabilities. This approach is theoretically equivalent to SS sampling in the sense that it computes identical Bayes Factors, at least when tree priors do not vary among the set of competed models (May and Rothfels, 2023). Reversible-jump MCMC has the advantage of accounting for uncertainty in model choice when estimating evolutionary parameters (i.e., tree topology, etc.), can accurately test alternative models involving variation in tree priors (May and Rothfels, 2023), and is far more computationally efficient than SS sampling, which requires many replicates. Nevertheless, it faces similar challenges to SS sampling regarding practical limits to the number of scientifically interesting models evaluated.

Regardless of which approach is used for model selection, the results of a model-fitting procedure only provide information about the relative fit of a model. It does not necessarily imply the model is adequately capturing the dynamics of the biological system being investigated. In fact, a best-fitting character model from a statistical standpoint of relative fit may poorly describe the underlying data. Determining whether a given model provides a reasonable explanation for data involves assessing a model’s adequacy. A variety of approaches to assessing model adequacy, including posterior predictive simulation (Höhna et al., 2018; Mulvey et al., 2024), are available for evaluating phylogenetic models of character evolution. We recommend researchers explore these newly developed tools to carefully interpret the implications and limits of their data.

Rather than exhaustively evaluate all possible alternative character models, investigate their adequacy, or infer the one “true” tree of olenid trilobites, we have focused on the far more modest goal of assessing whether different model-based assumptions about character evolution impact phylogenetic and macroevolutionary inferences using a dataset representative of many empirical studies. This is an important practical question, as the computational and analytical demands of model comparison is time consuming and a potential barrier to researchers who would prefer spending their time using phylogenies to solve taxonomic problems, describe new taxa, or investigate evolutionary patterns in their favorite clade. As systematic (palaeo)biologists ourselves, we fully empathize with this sentiment occasionally expressed by colleagues.

However, our results suggest inferences of phylogenetic relationships are highly sensitive to our assumptions about characters and how they evolved, and are not generally independent of other aspects of reconstructing a clade’s macroevolutionary history. We find that character evolution models have a significant impact on the probability of alternative tree topologies and influence which regions of treespace are explored. We recommend researchers think carefully about character evolution not only when interested in testing evolutionary hypotheses, but also when their primary research question solely revolves around inferring phylogenetic relationships.

## Conclusions

As models for the evolutionary “Tree of Life”, phylogenies form the cornerstone of evolutionary biology. When combined with quantitative models of character evolution, they provide the foundational framework for documenting the tempo and mode of phenotypic diversification. Because phylogenies for fossil taxa are estimated using morphological data, the generation of phylogenetic hypotheses using computational approaches are inextricably linked to models of morphological character evolution. Our results provide clear evidence that our assumptions about character evolution impact both the distribution of tree topologies inferred and macroevolutionary inferences derived from them, with the degree of model complexity and choice of character types (e.g., discrete vs. continuous traits) each having a significant influence on trees sampled during MCMC. As phylogenetic palaeobiology progress as a discipline, additional work is needed to refine our understanding of how alternative character evolution models may influence phylogenetic inferences more generally, especially for continuous data, which have received relatively little attention in treespace analyses compared to discrete-only datasets. We encourage palaeontological systematists to explore exciting new quantitative phylogenetic approaches in our field, but as always, be mindful of how assumptions about character evolution may shape our understanding of the Tree of Life.

## Supplementary material

Supporting information (Appendix S1) contains text further describing the construction and evaluation of tree spaces, all MCC trees and discussion on their potential implications for olenid systematics, and all supplemental figures. Appendix S1 is available online at: [link goes here]

## Data and code availability statement

Morphological data and scripts have been deposited in the Dryad Digital Repository (https://doi.org/xx.xxxx)

## Acknowledgements

We thank Jennifer Hoyal Cuthill, Sally Thomas, Kenneth De Beats, Selina Cole, and two anonymous reviewers for reading the manuscript and providing comments to improve its content and clarity. D.F.W. thanks Kurt Vonnegut for the concept of the chronosynclastic infundibulum, which inspired his perspectives about Bayesian approaches to simultaneous inference in phylogenetic palaeobiology.

## Author contributions

**Conceptualization** DF Wright (DFW), **Data curation** DFW, MJ Hopkins (MJH); **Formal analysis** DFW, MJH; **Investigation** DFW, MJH; **Methodology** DFW, MJH; **Visualization** DFW, MJH; **Writing – original draft** DFW; **Writing – review & editing** DFW, MJH

